## Supplementary materials for "Fine-scale family structure shapes influenza transmission risk in households: insights from a study of primary school students in Matsumoto city, 2014/15"

### 1. Model selection

#### Model selection on the complexity

In the first round of model selection, we compared models with different complexity. Our household transmission model was mainly characterised by two components, the effective household contact $\eta_{kl} = \beta \frac{c_{kl}}{C_k{}^{\gamma}}$ and the risk of external infection $\varepsilon_k$. Models corresponding to all possible combinations of assumptions were compared based on the Widely-applicable Bayesian information criterion (WBIC) (1). WBIC has the same scale as Bayesian information criterion. A difference of 2 in WBIC is considered as an indication of statistical significance, while a difference greater than 5 is deemed as a strong support. Table S1 compares the candidate models and their WBIC.

$c_{kl}$ and $\varepsilon_k$ were estimated as a single parameter $c_{kl} = c$ and $\varepsilon_k = \varepsilon$ under "Homogeneous"/"Uniform" assumptions, respectively. $\gamma$ was fixed at 0 in "DD" (density-dependent) models and 1 in "FD" (frequency-dependent), and was freely estimated in "IM" (intermediate) models. In "Single parent-Y" models, fathers and mothers who do not live with a spouse were classified as an additional type "single parent" (thus the number of types was 6 in these models). The best model (Model 12) was selected with a very strong support: ΔWBIC from the second best model was 16.9.

#### Model selection on the contact pattern matrix

After selecting the model complexity, we further tried to explore different contact pattern matrices $c_{kl}$. Let the rows and columns of $c_{kl}$ correspond to (Student, Sibling, Father, Mother, Other). Five parameters ($c_{\mathrm{CC}}, c_{\mathrm{FC}}, c_{\mathrm{MC}}, c_{\mathrm{OC}}, c_{\mathrm{AA}}$) being denoted by numbers 1 to 5, the contact pattern matrix $c_{kl}$ in the previous model selection had the following structure:

$$c_{kl} = \begin{bmatrix} 1 & 1 & 2 & 3 & 4 \\ 1 & 1 & 2 & 3 & 4 \\ 2 & 2 & 5 & 5 & 5 \\ 3 & 3 & 5 & 5 & 5 \\ 4 & 4 & 5 & 5 & 5 \end{bmatrix} \quad \text{(S1)}$$

Note that the diagonal elements for student, father and mother are displayed only for completeness and not used in the analysis (households in the dataset do not contain more than one students/fathers/mothers). Parameter estimates in Model 12 are shown in Table S2. In this contact pattern matrix, as all adults are assumed to share the same contact intensity. Meanwhile, the estimates of $c_{\mathrm{FC}}$ and $c_{\mathrm{OC}}$ are relatively similar. We explored variant models that further stratify $c_{\mathrm{AA}}$ while

$c_{\mathrm{FC}}$ and $c_{\mathrm{OC}}$ are equated to keep the number of parameters unchanged (=5).

We considered the following submodels: Model 12a (intense contact within couples), Model 12b (mother acting as a hub) and Model 12c (generation-assortative).

$$c_{kl}\text{ (Model 12a)} = \begin{bmatrix} 1 & 1 & 2 & 3 & 2 \\ 1 & 1 & 2 & 3 & 2 \\ 2 & 2 & 4 & 4 & 5 \\ 3 & 3 & 4 & 4 & 5 \\ 2 & 2 & 5 & 5 & 5 \end{bmatrix},$$

$$c_{kl}\text{(Model 12b)} = \begin{bmatrix} 1 & 1 & 2 & 3 & 2 \\ 1 & 1 & 2 & 3 & 2 \\ 2 & 2 & 5 & 4 & 5 \\ 3 & 3 & 4 & 4 & 4 \\ 2 & 2 & 5 & 4 & 5 \end{bmatrix}, \qquad \text{(S2)}$$

$$c_{kl}\text{(Model 12c)} = \begin{bmatrix} 1 & 1 & 5 & 3 & 5 \\ 1 & 1 & 5 & 3 & 5 \\ 5 & 5 & 2 & 2 & 5 \\ 3 & 3 & 2 & 2 & 5 \\ 5 & 5 & 5 & 5 & 4 \end{bmatrix}$$

Estimated contact pattern matrices are shown in Tables S3-S5. Models 12a and 12c had much better WBIC than Model 12 (ΔWBIC = -14.4 and ΔWBIC = -17.3, respectively), while that of Model 12b was slightly worse than Model 12. Of the two models exhibiting improved WBICs, Model 12c was selected with a significant WBIC difference of 2.9. Parameter estimates other than $c_{kl}$ did not vary between compared models to the first significant figure.

**Selection of the scaling factor**

In our baseline model, total amount of contacts $C_k = \sum_l c_{kl}$ was used to scale the effective household contact (i.e., $\eta_{kl} \propto {C_k}^{-\gamma}$) to reflect heterogeneous contact patterns. On the other hand, previous modelling studies often used household size $N$ in place of $C_k$ (2–5). Although $C_k$ and $N$ are correlated ($C_k$ and $N$-1 coincide in homogeneous settings) and may work as a good proxy with each other, we considered comparison between these two approaches to be of interest. We tested a variant of Model 12c where $C_k$ is replaced with $N$ (i.e., $\eta_{kl} \propto N^{-\gamma}$), but the model performance was significantly worsened (ΔWBIC = 8.8). The estimated value of gamma did not change ($\gamma$ = 0.52; CrI: 0.34-0.75). The use of total amount of contacts is preferred to household size as a scaling factor for the within-household transmission, and even when household size is used as variable, the semi-density-dependent model may still be applicable.

Table S1. WBIC of models with different sets of assumptions.

| Model ID | $c_{kl}$ | $\varepsilon_k$ | $\gamma$ | Single parent | WBIC | ΔWBIC |
|---|---|---|---|---|---|---|
| 1 | Hom | Unif | DD | N | 33269.16 | 2134.96 |
| 2 | Het | Unif | DD | N | 33054.70 | 1920.50 |
| 3 | Hom | Unif | FD | N | 33259.36 | 2125.16 |
| 4 | Het | Unif | FD | N | 32731.10 | 1596.90 |
| 5 | Hom | Unif | IM | N | 33243.32 | 2109.12 |
| 6 | Het | Unif | IM | N | 32277.92 | 1143.72 |
| 7 | Hom | Strat | DD | N | 31215.16 | 80.96 |
| 8 | Het | Strat | DD | N | 31151.08 | 16.88 |
| 9 | Hom | Strat | FD | N | 31205.44 | 71.24 |
| 10 | Het | Strat | FD | N | 31150.78 | 16.58 |
| 11 | Hom | Strat | IM | N | 31186.64 | 52.44 |
| 12 | Het | Strat | IM | N | 31134.20 | 0 |
| 13 | Hom | Unif | DD | Y | 33267.72 | 2133.52 |
| 14 | Het | Unif | DD | Y | 33061.14 | 1926.94 |
| 15 | Hom | Unif | FD | Y | 33256.72 | 2122.52 |
| 16 | Het | Unif | FD | Y | 32752.26 | 1618.06 |
| 17 | Hom | Unif | IM | Y | 33241.46 | 2107.26 |
| 18 | Het | Unif | IM | Y | 32182.18 | 1047.98 |
| 19 | Hom | Strat | DD | Y | 31223.68 | 89.48 |
| 20 | Het | Strat | DD | Y | 31167.16 | 32.96 |
| 21 | Hom | Strat | FD | Y | 31212.52 | 78.32 |
| 22 | Het | Strat | FD | Y | 31168.08 | 33.88 |
| 23 | Hom | Strat | IM | Y | 31194.82 | 60.62 |
| 24 | Het | Strat | IM | Y | 31151.98 | 17.78 |

Hom: homogeneous mixing, Het: Heterogeneous mixing

Unif: uniform risk of external infection, Strat: stratified risk of external infection

DD: density-dependent, FD: frequency-dependent, IM: intermediate

Single-Parent: whether a single parent have a unique parameter. Y=Yes, N=No.

WBIC: Widely-applicable Bayesian information criterion, ΔWBIC: WBIC difference from the best model

Table S2. Estimated contact pattern matrix ($c_{kl}$) in Model 12.

<table>
<tr><th></th><th>Student</th><th>Sibling</th><th>Father</th><th>Mother</th><th>Other</th></tr>
<tr><td>Student</td><td rowspan="2" colspan="2">1.28</td><td rowspan="2">0.54</td><td rowspan="2">1.40</td><td rowspan="2">0.45</td></tr>
<tr><td>Sibling</td></tr>
<tr><td>Father</td><td colspan="2">0.54</td><td rowspan="3" colspan="3">1</td></tr>
<tr><td>Mother</td><td colspan="2">1.40</td></tr>
<tr><td>Other</td><td colspan="2">0.45</td></tr>
</table>

WBIC = 31134.20; ΔWBIC = 0 (baseline)

Table S3. Estimated contact pattern matrix ($c_{kl}$) in Model 12a.

<table>
<tr><th></th><th>Student</th><th>Sibling</th><th>Father</th><th>Mother</th><th>Other</th></tr>
<tr><td>Student</td><td rowspan="2" colspan="2">0.97</td><td rowspan="2">0.39</td><td rowspan="2">1.09</td><td rowspan="2">0.39</td></tr>
<tr><td>Sibling</td></tr>
<tr><td>Father</td><td colspan="2">0.39</td><td rowspan="2" colspan="2">1</td><td rowspan="2">0.39</td></tr>
<tr><td>Mother</td><td colspan="2">1.09</td></tr>
<tr><td>Other</td><td colspan="2">0.39</td><td colspan="2">0.39</td><td>1</td></tr>
</table>

WBIC = 31119.78; ΔWBIC = -14.42

Table S4. Estimated contact pattern matrix ($c_{kl}$) in Model 12b.

<table>
<tr><th></th><th>Student</th><th>Sibling</th><th>Father</th><th>Mother</th><th>Other</th></tr>
<tr><td>Student</td><td rowspan="2" colspan="2">1.25</td><td rowspan="2">0.49</td><td rowspan="2">1.37</td><td rowspan="2">0.49</td></tr>
<tr><td>Sibling</td></tr>
<tr><td>Father</td><td colspan="2">0.49</td><td>1.01</td><td rowspan="3">1</td><td>1.01</td></tr>
<tr><td>Mother</td><td colspan="2">1.37</td><td></td><td></td></tr>
<tr><td>Other</td><td colspan="2">0.49</td><td>1.01</td><td>1.01</td></tr>
</table>

WBIC = 31134.72; ΔWBIC = 0.54

Table S5. Estimated contact pattern matrix ($c_{kl}$) in Model 12c.

<table>
<tr><th></th><th>Student</th><th>Sibling</th><th>Father</th><th>Mother</th><th>Other</th></tr>
<tr><td>Student</td><td rowspan="2" colspan="2">1.04</td><td rowspan="2">0.43</td><td rowspan="2">1.16</td><td rowspan="4">0.43</td></tr>
<tr><td>Sibling</td></tr>
<tr><td>Father</td><td colspan="2">0.43</td><td rowspan="2" colspan="2">1</td></tr>
<tr><td>Mother</td><td colspan="2">1.16</td></tr>
<tr><td>Other</td><td colspan="4">0.43</td><td>1.97</td></tr>
</table>

WBIC = 31116.88; ΔWBIC = -17.32 (best model)

### 2. Source-stratified risk of infection and risk attributable to introduction of influenza into a household

We quantified the risk of infection attributable to external and within-household infection from the parameter estimates. Three family compositions were selected as model cases: (a) "nuclear family": father, mother and two children, (b) "many-siblings family": father, mother and four children, and (c) "three-generation family": father, mother, two children and two grandparents. We assumed that one of the children in each model case households was "student", and the others were "siblings". Overall risk of infection for type $k$ individual is given by

$$r_k = \sum_{\boldsymbol{n}} \frac{n_k}{N_k} \pi(\boldsymbol{n}; \boldsymbol{N}, \boldsymbol{\varepsilon}, H), \tag{S3}$$

(the sum is taken for all possible $\boldsymbol{n}$), and $r_k - \varepsilon_k$ corresponds to the additional infection risk due to within-household transmission.

We also compared the risk of infection after the introduction of influenza into households with the initial overall risk. We defined post-introduction risk as the conditional probability that an individual experience infection by the end of season, given that one index case is already observed in the same family. Here, for simplicity, we limited the analysis to introductions by primary school students (i.e., individual type "student") only.

Suppose that $k$=1 corresponds to the type "student". Post-introduction risk obtained by modifying the formula for $r_k$ as

$$r_k^{\text{pos}} = \sum_{\{\boldsymbol{n}|n_1=0\}} \frac{n_k}{N_k} \cdot \frac{\pi(\boldsymbol{n}; \boldsymbol{N}, \boldsymbol{\varepsilon} + \boldsymbol{H}_1, H)}{S_1(\boldsymbol{n}, \boldsymbol{\varepsilon})}. \tag{S4}$$

$\boldsymbol{H}_1$ is the (additional) force of infection arising from the infected student, i.e., $(\boldsymbol{H}_1)_k = \frac{c_{k1}}{{C_k}^{\gamma}}$. The sum is taken for all possible $\boldsymbol{n}$ whose first component $n_1 = 0$ (because the force of infection from the student is incorporated in $\boldsymbol{H}_1$). Note that by dividing $h(\boldsymbol{n}; \boldsymbol{N}, \boldsymbol{\varepsilon} + \boldsymbol{H}_1, H)$ by $S_1(\boldsymbol{n}, \boldsymbol{\varepsilon})$, we can yield the conditional probability that $\boldsymbol{n}$ individuals (other than the "student") are infected given the presence of the force of infection $\boldsymbol{H}_1$.

### 3. Sensitivity analysis

#### Procedures for the sensitivity analysis

(i) Ascertainment bias

In order to account for potential ascertainment bias, we incorporated reporting probabilities into the model. We assumed that infections are reported with a certain probability $p_k$. Epidemiological properties, such as infectiousness, were assumed to be identical between reported and unreported cases. The likelihood of observing a household final size outcome $(\boldsymbol{n}; \boldsymbol{N})$ given the reporting probability vector $\boldsymbol{p}$ is obtained by using the binomial distribution:

$$L(\boldsymbol{\varepsilon}, H, \boldsymbol{p}; (\boldsymbol{n}; \boldsymbol{N})) = \sum_{\boldsymbol{n}' \geq \boldsymbol{n}} h(\boldsymbol{n}'; \boldsymbol{N}, \boldsymbol{\varepsilon}, H) \prod_k \mathrm{Bin}(n_k; n'_k, p_k). \tag{S5}$$

The sum $\sum_{\boldsymbol{n}' \geq \boldsymbol{n}}$ is taken for all vector ***n'*** satisfying $n_k \leq n_k{}' \leq N_k$ $(\forall k)$.

In this sensitivity analysis, we assumed that the reporting probability $p$ for children ("student" and "sibling") is 0.8. The reporting probability for adults was varied from 0.5 to 0.8.

(ii) Different susceptibility in children

Susceptibility to influenza infection was differentiated between children adults. Let $\sigma$ be the susceptibility of children relative to that of adults. The effect of $\sigma$ was employed in the model by differentiating the transmissibility $\beta$ as

$$\beta_k = \begin{cases} \beta\sigma & (k = \text{"Student"}, \text{"Sibling"}) \\ \beta & (\text{otherwise}) \end{cases}. \tag{S6}$$

Five different values of $\sigma$: 0.75, 1.25, 1.5, 1.75, 2.0 were tested. $\sigma$ = 0.75 corresponds to the assumption that children may have less risk of infection per exposure (e.g., due to potentially high vaccination coverage). $\sigma$ greater than 1 reflects the assumption that children are more vulnerable than adults.

(iii) Multiple counting of households

We identified all the possible combinations of respondents who might be from the same household by the following process. (1) Respondents were classified by their school and family composition. We assumed that siblings usually go to the same primary school. (2) Data were matched up, and consistency was checked between the sex and grade of the respondent and the reported composition of siblings. For instance, a second-grade boy who has no older brother should not be from the same household as a fourth-grade girl who has an older brother (the girl's older brother should also be the boy's older brother). Here, we assumed that siblings should be in different grades, and neglected the possibility of twins or siblings in the same grade. (3) Individuals potentially from the same household were grouped together. Combination for grouping was chosen so that as many individuals as possible are grouped together in total. Individuals in each matched group were assumed to be from the same household, and their data were integrated to represent one household data. Respondents were classified as "students", and siblings who were not found in the dataset was classified as "siblings". Because sex, school and grade were not used in the parameter estimation, individual-level details of the grouping arrangement (who was grouped with whom) did not affect the subsequent analysis. Through this whole process, 1,294 individuals identified as candidates potentially from multiple-counted households were processed, reducing the number of households from 10,486 to 9,763 (-6.9%). Note that this is an extreme case where as many consistent siblings as possible are grouped together, and that the reality may lie between the two extremes (no-grouping and maximum-grouping).

(iv) Case censoring

Participants reported in the survey the total number of siblings, siblings in four categories (older brother, older sister, younger brother or younger sister) and whether siblings in each category had influenza. Let $\boldsymbol{M} = (M_1, M_2, M_3, M_4)$ and $\boldsymbol{m} = (m_1, m_2, m_3, m_4)$ be the true composition of siblings and the number of siblings with an influenza episode in each category (1: older brother; 2: older sister, 3: younger brother; 4: younger sister), respectively. Due to the questions in the survey, the dataset did not include either $\boldsymbol{M}$ or $\boldsymbol{m}$. Instead we have $M = \sum_i M_i$ and censored sibling data $\boldsymbol{M}'$ ($M_i' = \min(M_i, 1)$) and $\boldsymbol{m}'$ ($m_i' = \min(m_i, 1)$), as the questions on sibling categories were yes-no questions.

We constructed a modified likelihood function to address this censoring issue. The basic idea was to generate all possible patterns of $\boldsymbol{M}$ and $\boldsymbol{m}$ that are consistent with the observation and aggregate the corresponding probabilities to obtain the likelihood for the censored data. First, we defined a conditional probability $\pi(\boldsymbol{M}; M)$, the probability that the true sibling composition is $\boldsymbol{M}$ given $M$. Assuming that the probability of being the $n$-th child in given $M$ siblings is equally $\frac{1}{M+1}$ and that the sex of a child is evenly distributed, we get

$$\pi(\boldsymbol{M}; M) = \frac{1}{(M+1)\cdot 2^M}\binom{M_1 + M_2}{M_1}\binom{M_3 + M_4}{M_3}\sigma(\boldsymbol{M}, M), \tag{S7}$$

where $\sigma(\boldsymbol{M}, M)$ is an indicator function that takes 1 if $\boldsymbol{M}$ is consistent with $M$ (i.e., $\sum_i M_i = M$), and 0 otherwise.

Let $\pi(\boldsymbol{m}; \boldsymbol{M}, \varphi)$ be the probability of observing a sibling outcome pattern $\boldsymbol{m}$ given $\boldsymbol{M}$. This is also conditional to the existence of other family members and their outcomes, and those conditions are represented by $\varphi$.

Using $\pi(\boldsymbol{M}; M)$ and $\pi(\boldsymbol{m}; \boldsymbol{M}, \varphi)$, the likelihood of observing $\{\boldsymbol{M}', \boldsymbol{m}'\}$ given $M$ and $\varphi$ is

$$l(\boldsymbol{M}', \boldsymbol{m}'; M, \varphi) = \sum_{\boldsymbol{M}} \pi(\boldsymbol{M}; M)\,\sigma(\boldsymbol{M}, \boldsymbol{M}') \sum_{\boldsymbol{m}} \pi(\boldsymbol{m}; \boldsymbol{M}, \varphi)\,\sigma(\boldsymbol{m}, \boldsymbol{m}'), \tag{S8}$$

where $\sigma(\boldsymbol{M}, \boldsymbol{M}')$ and $\sigma(\boldsymbol{m}, \boldsymbol{m}')$ are indicator functions checking if $\boldsymbol{m}$ and $\boldsymbol{M}$ are consistent with the observation.

Since we assume all siblings exhibit identical epidemiological behaviour, considering the effect of loss of distinguishability, $\pi(\boldsymbol{m}; \boldsymbol{M}, \varphi)$ is substituted with

$$\pi(\boldsymbol{m}; \boldsymbol{M}, \varphi) = \frac{\prod_i \binom{M_i}{m_i}}{\binom{M}{m}}\pi(m; M, \varphi), \tag{S9}$$

where $m = \sum_i m_i$. $\pi(m; M, \varphi)$ is equivalent to $\pi$ in equation (1) in the main text, and thereby we get the likelihood accounting for possible case censoring in siblings.

**Results of the sensitivity analysis**

The estimates from the sensitivity analysis were compared in Figure S1. Equally lowering the reporting probability for children and adults slightly increased some of the parameters while the overall relative magnitude was almost conserved. When the reporting probability for adults was set lower than children, parameters which involve adults increased and those involving children decreased (Figures S1A and S1B). Increasing the relative susceptibility in children resulted in lower child-involved contact intensities (Figures S1C and S1D). Multiple counting of data reported by students from the same household did not seem to have affected the result, but some changes were caused by addressing censored cases in siblings, which may be resulted from the possibility of unobserved sibling cases (Figures S1E and S1F). Except that the contact intensity between children was substantially lowered by either underreporting of adults or high susceptibility in children, the relative trend remained almost similar throughout our sensitivity analysis. Especially, the risk of external infection in children and the contact intensity between children and adults remained at the sufficient level, such that the secondary transmission from children is still of paramount importance.

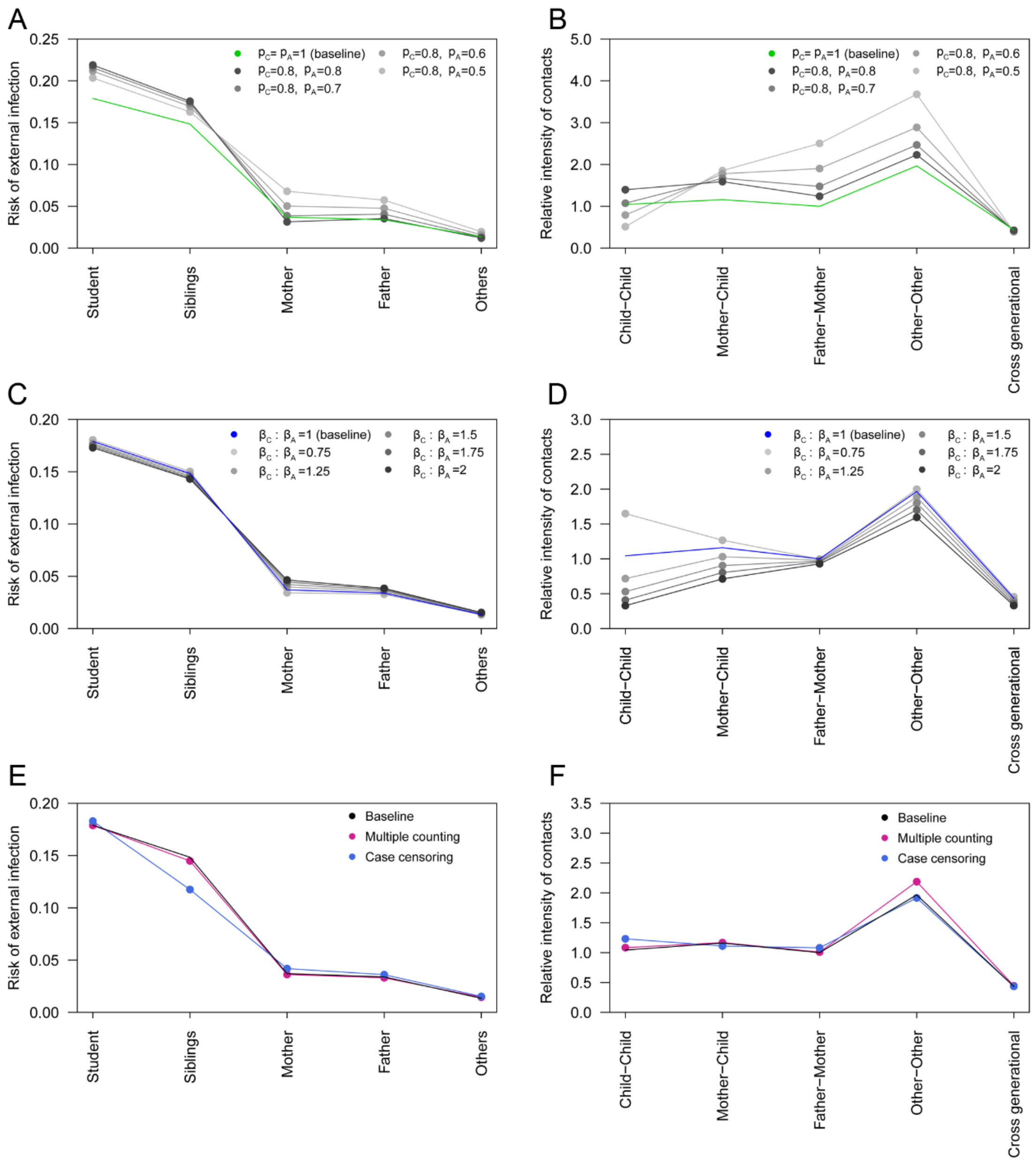


Figure S1. Parameter estimates from the sensitivity analysis. The estimated risk of external infection and relative intensity of household contacts are compared with the baseline estimates. Relative intensity of contacts in the figures is multiplied by the relative change in the estimated transmissibility parameter $\beta$ for comparability.

(A), (B) Various reporting probabilities in children ($p_C$) and adults ($p_A$).

(C), (D) Various ratios between susceptibility in children ($\beta_C$) and adults ($\beta_A$).

(E), (F) Estimates from the modified dataset addressing multiple counting of households and censoring of cases in siblings.

### 4. Model fit

To evaluate the goodness-of-fit of our model, the model prediction was compared with the observed data. Let $\hat{\theta}$ be the set of median parameter estimates. $\pi(\boldsymbol{n};\boldsymbol{N},\hat{\theta})$, the probability of observing outcome pattern $\boldsymbol{n}$ given household composition $\boldsymbol{N}$, is obtained from Equation (1) in the main text. Assuming that the distribution of $\boldsymbol{N}$ in dataset $D$ is given as observed ($\pi_D(\boldsymbol{N})$), the predictive distribution of the outcome patterns $(\boldsymbol{N}_i,\boldsymbol{n}_i)$ (approximated by the point estimate $\hat{\theta}$) is

$$\pi(\boldsymbol{N}_i,\boldsymbol{n}_i;\hat{\theta}) = \pi(\boldsymbol{n}_i;\boldsymbol{N}_i,\hat{\theta})\pi_D(\boldsymbol{N}_i), \tag{S10}$$

Figure S2 compares the predictive distribution with the actual frequency in the dataset. The 95% intervals are approximated by the 95% quantiles of a binomial distribution

$$F_D(\boldsymbol{N},\boldsymbol{n}) \sim \mathrm{Binom}\left(F_D(\boldsymbol{N}),\pi(\boldsymbol{n};\boldsymbol{N},\hat{\theta})\right), \tag{S11}$$

where $F_D$ is the frequency in data $D$ of size $S_D$. The predicted and observed frequency show a good accordance despite the relatively modest parameter space dimension (=11). The similarity between the two distributions are also supported by the empirical Kullback-Leibler divergence of 0.05, where

$$\widehat{\mathrm{KL}} = \sum_d \frac{F_D(\boldsymbol{N},\boldsymbol{n})}{S_D}\cdot\log\left(\frac{F_D(\boldsymbol{N},\boldsymbol{n})}{\pi(\boldsymbol{n};\boldsymbol{N},\hat{\theta})F_D(\boldsymbol{N})}\right). \tag{S12}$$

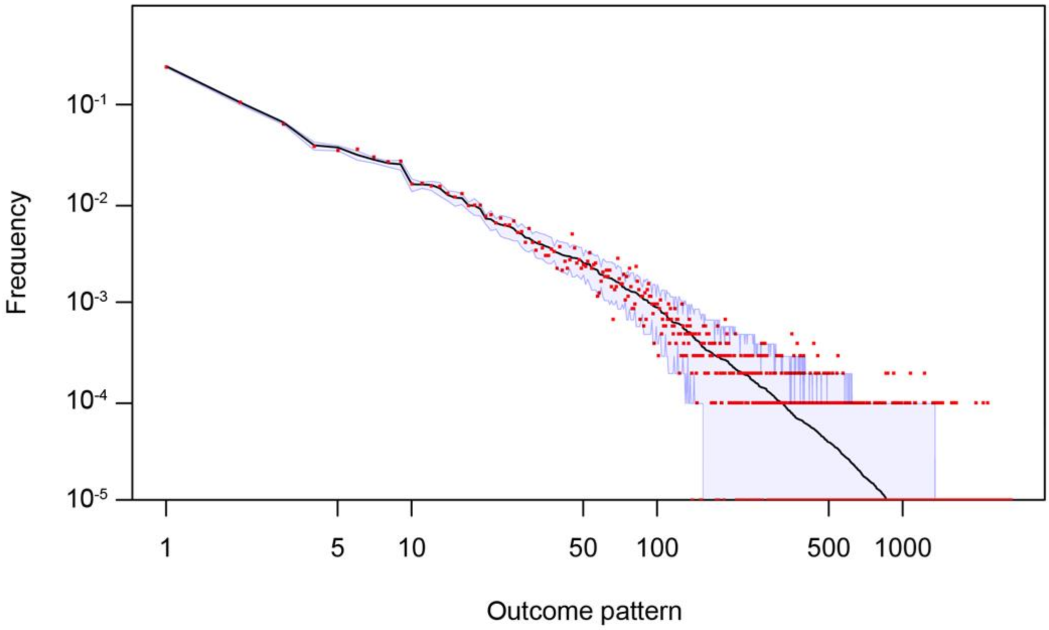


Figure S2. Comparison between the predicted and observed household final outcomes.

Red dots correspond to the observed relative frequency of data (household compositions and final outcomes of the household members), where the x-axis denotes the numbering of outcome patterns $(\boldsymbol{N},\boldsymbol{n})$. With the sample size of ~10,000, $10^{-4}$ on the y-axis denotes frequency 1; dots for frequency 0 are shown on the x-axis. The black line indicates the probability of observation predicted by the model, and the shaded area shows 95% intervals. Both x- and y-axes are on a logarithmic scale.
